# Dalbavancin and wound healing: new evidences/insights in a mouse model of skin infection

**DOI:** 10.1101/805895

**Authors:** Oriana Simonetti, Guendalina Lucarini, Gianluca Morroni, Fiorenza Orlando, Raffaella Lazzarini, Antonio Zizzi, Lucia Brescini, Mauro Provinciali, Andrea Giacometti, Annamaria Offidani, Oscar Cirioni

**Affiliations:** Clinic of Dermatology, Department of Clinical and Molecular Sciences, Polytechnic University of Marche, Ancona, Italy; Histology, Dept. of Clinical and Molecur Sciences, Polytechnic University of Marche, Ancona, Italy; Clinic of Infectious Diseases, Dept. of Biomedical Sciences and Public Health, Polytechnic University of Marche, Ancona, Italy; Experimental Animal Models for Aging Unit, Scientific Technological Area, I.N.R.C.A. I.R.R.C.S., Ancona, Italy; Pathologic Anatomy and Histopathology, Department of Biomedical Sciences and Public Health, Polytechnic University of Marche, Ancona, Italy.

**Author notes:** both the authors equally contributed to this work. **Correspondence to:** Oriana Simonetti, MD, PhD, Clinic of Dermatology, Department of Clinical and Molecular Sciences, Polytechnic University of Marche, c/o Ospedali Riuniti. Via Conca 71, 60126 Ancona AN, E.mail.

## Abstract

Dalbavancin is an effective antibiotic widely used to treat skin infection. Our aim was **t**o determinate the effects of dalbavancin administration on wound healing compared to vancomycin, and to elucidate if EGFR, MMP-1, MMP-9 and VEGF could be involved in its therapeutic mechanism.

A mouse model of MRSA skin infection was established. Mice were treated daily with vancomycin (10mg/kg) and weekly with dalbavancin, at day 1 (20 mg/kg) and day 8 (10 mg/kg). After 14 days wounds were excised and bacterial counts were perfomed. Wound healing was assessed by histological and immunohistochemical staining, followed by protein extraction and immunoblotting. Our microbiological results confirmed that both dalbanvancin and vancomycin are effective in reducing the bacterial load in wounds. Dalbavancin had a strong effect compared with infected untreated animals and vancomycin treated group. The wounds treated with dalbavancin showed robust epidermal coverage with a reconstitution of the regular and keratinized epidermal lining and a well-organized granulation tissue with numerous blood vessels, although slightly less than in the uninfected group, while in vancomycin treated group the epithelium appeared in general still hypertrophic, the granulation tissue appears even less organized.

We observed elevated EGFR and VEGF expression in both treated groups, although it was higher in dalbavancin treated mice. MMP-1 and MMP-9 were decreased in uninfected and in both treated tissue when compared with untreatd infected wounds.

This study showed faster healing with dalbavancin treatment that might be associated with a higher EGFR and VEGF levels.

## INTRODUCTION

It has recently proposed a new classification of skin and soft tissue infections namely acute bacterial skin and skin structure infections (ABSSSIs) where Superficial incisional Surgical Site Infections or SSIs (involving only skin or subcutaneous tissue of the incision) are recognized a subgroup ABSSSIs (1). In particular SSIs constitute a significant burden for the healthcare system, because of the associated morbidity, mortality, length of hospital stay, a higher probability of intensive care unit (ICU) admission with overall direct and indirect costs (2).

The common pathogens that are resposible of SSI are *Staphylococcus aureus*, followed by Coagulase-negative staphylococci and *Enterococcus* species. In particular methicillin-resistant *Staphylococcus aureus* (MRSA) is considered a leading cause of human nosocomial infections (3) as well as community-associated MRSA (CA-MRSA) is associated to deadly infection (4) and livestock-associated MRSA (LA-MRSA) represent major reservoir of infection (5).

In the last decades the epidemiological aspects of nosocomial infection as well as the growing prevalence of CA-MRSA have been increasingly studied to conteract this worldwide public health problem. Particular attention has been paid to the problem of LA-MRSA. Several studies demonstrated that this superbug can be trasmitted to human by either healthy and disease animals, usually cow, horses and companion animals (6–7).

Taking into account these considerations, the respect of the principles of a good antibiotic stewardship is mandatory for fighting antimicrobial resistance and aim, next to optimization of clinical outcomes, to the prevention of the spread of antibiotic-resistant strains (8). Dalbavancin is a novel antimicrobial agent, belonging to the lipoglycopeptides family (a subclass of glycopeptides), active against *S. aureus* (including MRSA), *Streptococcus pyogenes*, *Streptococcus agalactiae*, *Streptococcus anginosus* group, *Enterococcus faecalis* and *Enterococcus faecium* (including ampicillin-resistant isolates) and approved for the treatment of ABSSSIs caused by susceptible Gram-positive organisms (9). Moreover, it has a bactericidal effect and an extremely extended half-life that make it a promising alternative to conventional antibacterials such as vancomycin, that is considered the standard of care for treatment of gram-positive skin and soft-tissue infections (10).

Wound healing is a highly complex physiological process involving ordered events where growth factors are critically important for coordinating cell-cell and cell-matrix interactions during normal injury repair. In particular Epidermal keratinocytes (KCs) respond to injury by becoming hyperproliferative, migratory and pro-inflammatory in processes regulated by several factors such as epidermal growth factor receptor (EGFR), a 170-kDa transmembrane protein belonging to ErbB family receptors, which includes three other members (ErbB2/HER-2, ErbB3/HER-3, ErbB4/HER-4). EGFR signaling plays key roles in skin regeneration and wound healing by stimulating KCs proliferation (11). Among the many proteins, that are essential for the restoration of tissue integrity, the metalloproteinase family is one of the most important. Matrix metalloproteinases (MMPs) comprise a large family of extracellular proteases, which are up-regulated during wound healing by epidermal, dermal, fibroblast, or blood cells in mammals (12). MMPs contribute significantly in reorganization of the extracellular matrix for numerous physiological functions including morphogenesis, tissue remodeling and wound healing (13). In particular MMP-1 and MMP-9 reflect the correlation between high protease activity and poor healing outcomes (12).

An essential feature of normal wound repair is the formation of granulation tissue, i.e. fibrovascular tissue containing fibroblasts, collagen and blood vessels. Vascular endothelial growth factor (VEGF) has shown to be important in wound healing by promoting the early events in angiogenesis, particularly endothelial cell migration and proliferation (14). VEGF is produced by endothelial cells, keratinocytes, fibroblast, smooth muscle cells, platelets, neutrophils and macrophages^15^, and, although in normal skin is negligible, in injured skin is markedly up-regulated (14–16).

This study aimed to explore the therapeutic effects of dalbavancin on wound healing in an infected MRSA mouse model, using histological, immunohistochemial and western blot analysis, and to elucidate if EGFR, MMP-1, MMP-9 and VEGF could be involved in its therapeutic mechanisms.

## RESULTS

### Bacterial Counts

Overall, data analysis showed that inhibition of bacterial growth was achieved in all treated groups. In details, mean bacterial numbers in untreated group (7,51 x10^8^ ± 5,05 x10^7^ CFU/ml) were significantly higher than those recovered from all treatment groups (Fig. 1). Treatment with vancomycin reduced the mean bacterial numbers to 8,04 x10^6^ ± 7,96 × 10^6^ CFU/ml, while dalbavancin had a strong effect compared with control infected untreated animals and vancomycin treated group (8,71 × 10^5^ ± 9,02 × 10^5^ CFU/ml).

**Figure 1.**
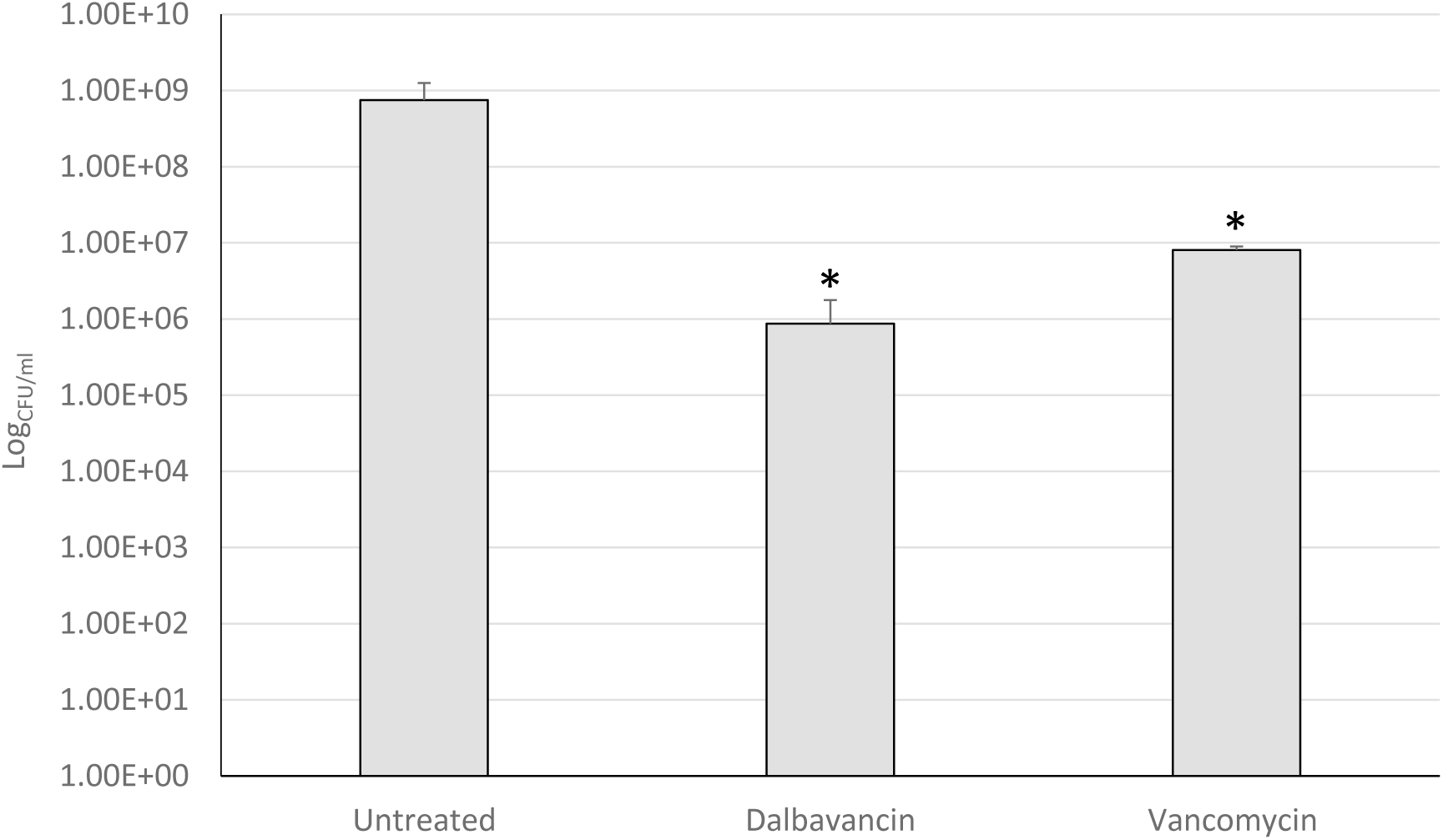
Bacterial counts of excised wound. Counts are expressed as log of CFU/ml. The limit of detection for this method was approximately 10 CFU/ml. * represent *p*<0.05 compared to infected but not treated group-

### Histological evaluation

To investigate the effects of the different treatments on wound healing, we analyzed the area of the wound, in particular reepithelialization, granulation tissue, collagen deposition and inflammatory cells according to wound healing scores^20^ summarized in Table 1. The statistical analysis was reported in Table 2 and the morphological features was shown in Figure 2.

**Table 1.**
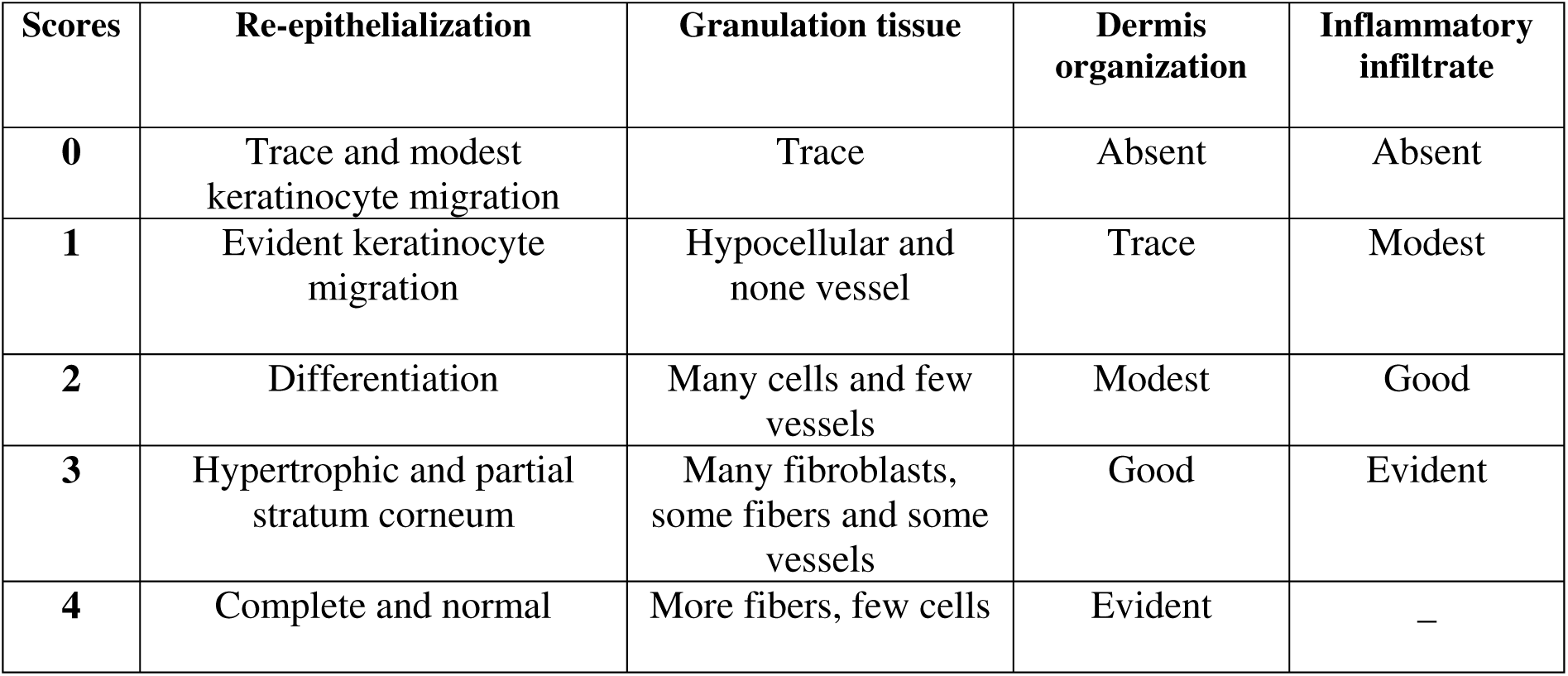
A five-tiered grading system to evaluate wound healing.

| **Scores** | **Re-epithelialization** | **Granulation tissue** | **Dermis organization** | **Inflammatory infiltrate** |
| --- | --- | --- | --- | --- |
| **0** | Trace and modest keratinocyte migration | Trace | Absent | Absent |
| **1** | Evident keratinocyte migration | Hypocellular and none vessel | Trace | Modest |
| **2** | Differentiation | Many cells and few vessels | Modest | Good |
| **3** | Hypertrophic and partial stratum corneum | Many fibroblasts, some fibers and some vessels | Good | Evident |
| **4** | Complete and normal | More fibers, few cells | Evident | – |

**Table 2.** Analysis of wound healing in the different mice groups according to the histological scores.

| **Model** | **Re-epithelialization**<br>**(mean±SD)** | **Granulation**<br>**tissue**<br>**(mean±SD)** | **Dermis**<br>**organization**<br>**(mean±SD)** | **Inflammatory**<br>**infiltrate**<br>**(mean±SD)** |
| --- | --- | --- | --- | --- |
| **Uninfected (C0)** | 3.85±0.25 | 3.52±0.22 | 3.85±0.50 | 1.02±0.02 |
| **Untreated infected**<br>**(C1)** | 0.50±0.08 | 1.25±0.08 | 1.80±0.30 | 2.00±0.25 |
| **Infected treated with**<br>**vancomycin (C2)** | 2.85±0.20 | 3.00±0.25 | 3.20±0.28 | 1.20±0.03 |
| **Infected treated with**<br>**dalbavancin (C3)** | 3.35±0.10 | 3.20±0.15 | 3.75±0.18 | 1.2±0.08 |
| **ANOVA test**<br>**Test di Bonferroni**<br>**p<0.05** | **C0 vs C1, C2, C3**<br>**C1 vs C0, C2, C3**<br>**C3 vs C2** | **C0 vs C1, C2**<br>**C1 vs C2,C3,** | **C0,C3 vs C0, C1**<br>**C2 vs C1** | **C1vs C0,C2,**<br>**C3** |

**Figure 2.**
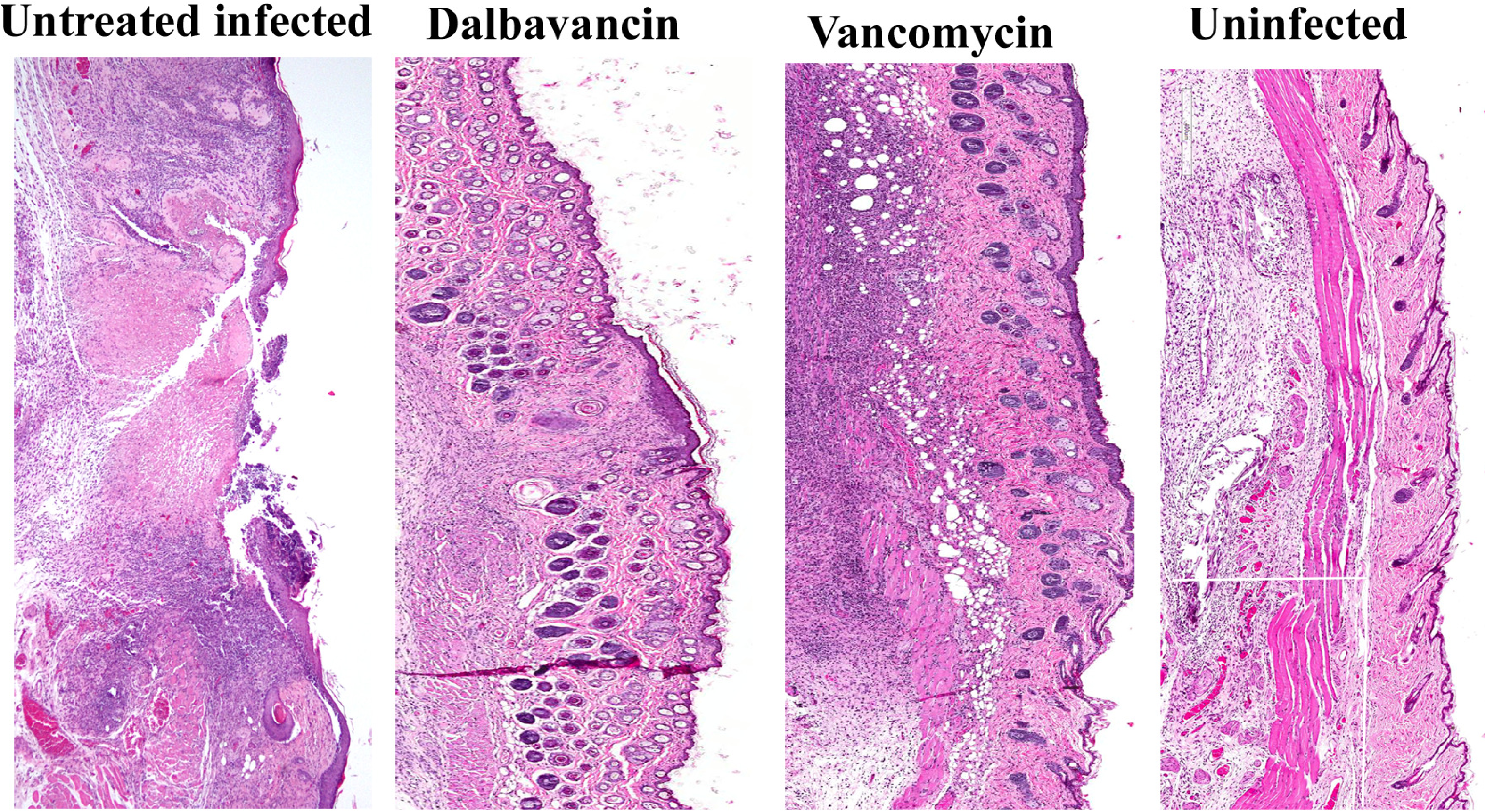
Representative images of hematoxylin- and eosin-stained histological sections of wound healing tissues from uninfected mice, infected rats treated with dalbavancin or vancomycin, and infected untreated mice on day 14 after injury. Wound healing was still incomplete in the untreated groups, both treated groups exhibited a good re-epithelialization; in particular, the wounds treated with Dalbavancin showed robust epidermal coverage and well organized granulation tissue (Original magnification 100x).

The extent of newly formed epithelium and collagen deposition, the increase in the thickness of granulation tissue and the decrease in the number of inflammatory cells were considered as progress in the healing process.

Wound healing was still incomplete in the untreated groups, as observed by the partial regeneration of the epidermis, poor organization of dermis layers, immature granulation tissue (many cells and some fibers), and abundant inflammatory infiltrate. These mice showed the worst degree of healing (p<0.05).

From a microscopic examination, the neo-epithelium developed significantly different patterns of regeneration among the treated groups: interestingly, the re-epithelialization of the wounds treated with dalbavancin showed robust epidermal coverage with a reconstitution of the regular and keratinized epidermal lining, comparable to the re-epithelialization of the uninfected mice, except in a small central area where the epithelium appears to be hypertrophic. The wounds treated with vancomycin showed a good re-epithelialization, although the epithelium appeared in general still hypertrophic when compared to dalbavancin and uninfected groups (p<0.05).

The wounds treated with dalbavancin showed a well-organized granulation tissue with numerous blood vessels, although slightly less than in the uninfected group. In the wounds treated with vancomycin the granulation tissue appears even less organized.

The treated wounds presented a distinct pattern of collagen distribution and remodeling similar to that of uninfected skin; in addition, mild or moderate inflammation was observed in the underlying connective tissue.

### Immunohistochemical evaluation

To investigate the influence of the dalbavancin and vancomycin treatment on the expression of VEGF, EGFR, MMP-9 and MMP-1 and in wound healing, an immunhistochemical evaluation was performed. The immunohistochemical staining is shown in Figures.3-4. and the data are summarized inTable 3.

**Figure 3.**
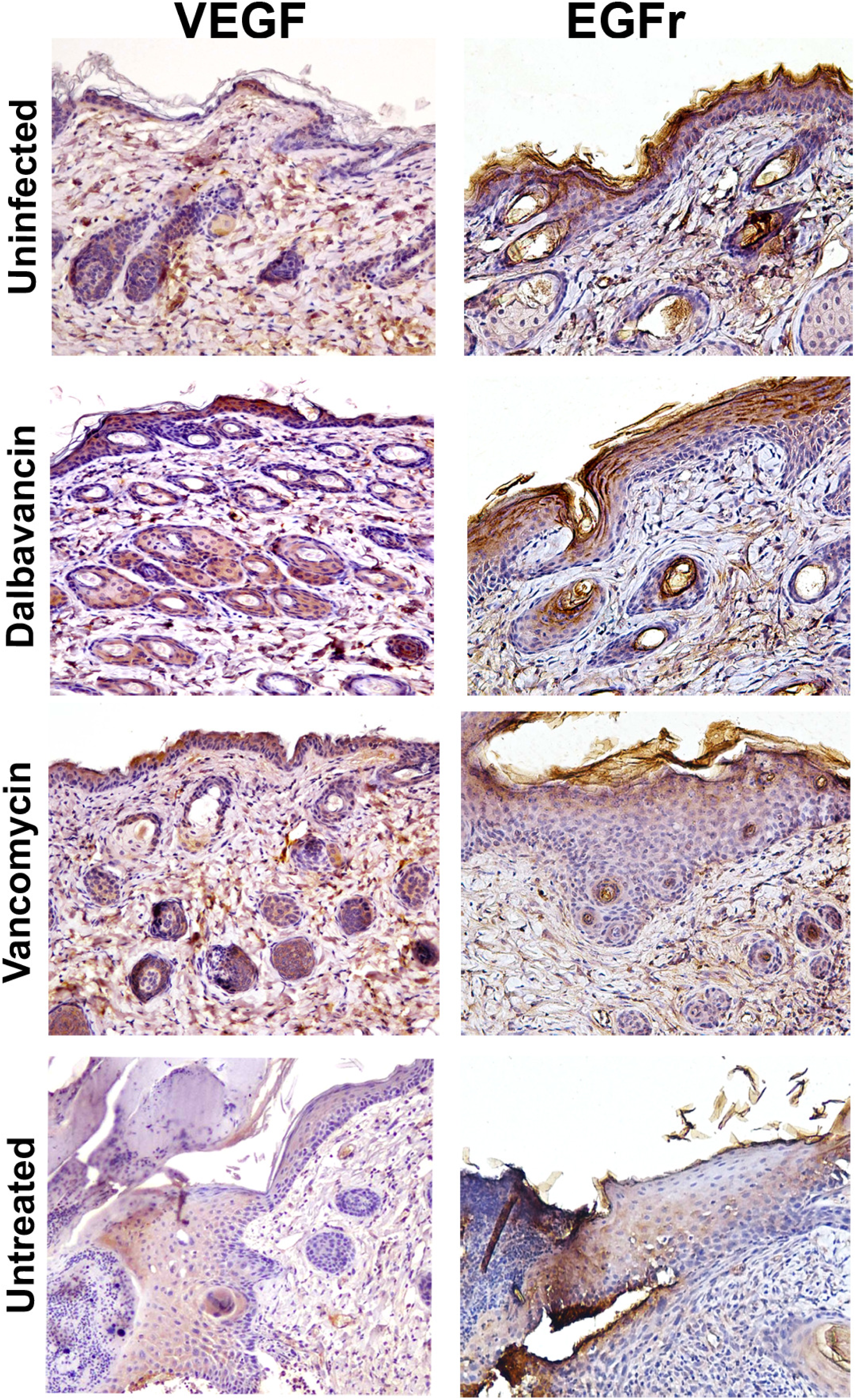
Immunohistochemical staining of VEGF and EGFr in wound healing tissues from uninfected mice, infected rats treated with dalbavancin or vancomycin, and infected untreated mice on day 14 after injury. VEGF and EGFr expression was greater in the uninfected and treated infected groups respect to the untreated, in particular the wounds treated with dalbavancin showed the highest value (immunoperoxidase; original magnification 200x).

**Figure 4.**
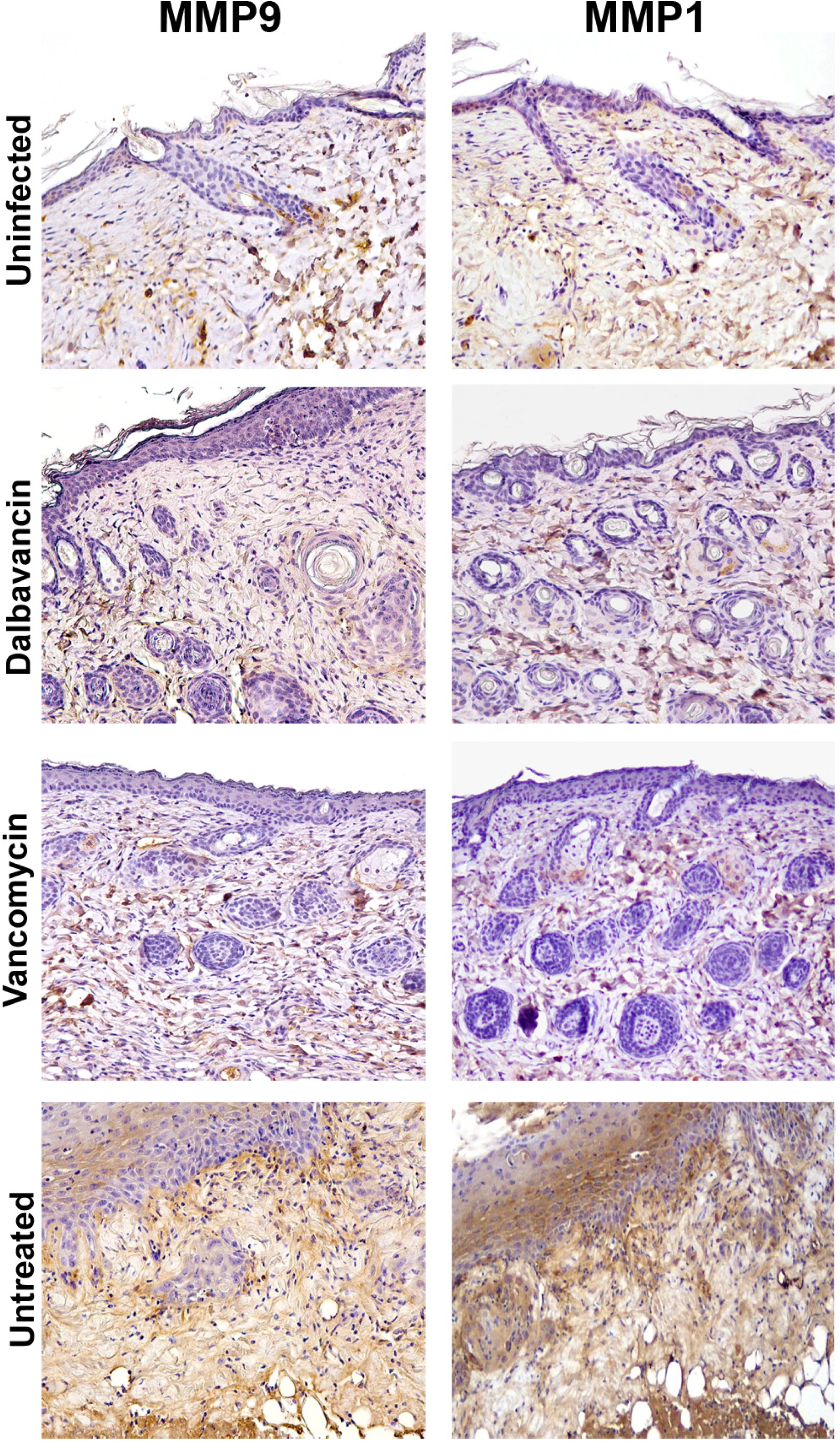
Immunohistochemical staining of MMP-9 and MMP-1 in wound healing tissues from uninfected mice, infected mice treated with dalbavancin or vancomycin, and infected untreated mice on day 14 after injury. MMP-9 and MMP-1 expression was high in the untreated infected wounds, the treated wounds showed the lowest values (immunoperoxidase; original magnification 200x).

**Table 3.** Immunohistochemical expression of VEGF, MMP2, MMP9, MMP1 and EGFr in the different mice groups.

| **Model** | **VEGF**<br>**(mean±SD)\*\*** | **EGFr**<br>**(mean±SD)** | **MMP-9**<br>**(mean±SD)** | **MMP-1**<br>**(mean±SD)** |
| --- | --- | --- | --- | --- |
| **Uninfected (C0)** | 30.0±3.2 | 40.0±2.5 | 20.3±2.6 | 22.0±1.2 |
| **Untreated Infected (C1)** | 15.5±2.6* | 31.0±2.5 | 37.0±1.6 | 45.0±5.2 |
| **Infected treated with vancomycin (C2)** | 31.6±3.2 | 30.0±2.0 | 16.2±1.8* | 18.0±3.6* |
| **Infected treated with dalbavancin (C3)** | 40.0±4.2* | 50.0±2.2* | 15.0±2.5* | 20.0±3.0* |
| **ANOVA test**<br>**Test di Bonferroni**<br>**p<0.05\*** | **C3 vs C0, C1, C2,**<br>**C1 vs C0, C2** | **C3 vs C0, C1, C2**<br>**C0, C3 vs C1, C2**<br>**C3 vs C0** | **C1 vs C0, C2, C3**<br>**C2,C3 vs C0** | **C1 vs C0, C2, C3** |

\*\* percentage of positive cells

VEGF expression was greater in the uninfected and treated infected groups respect to the untreated (p<0.05), the wounds treated with dalbavancin showed the highest VEGF values.

EGFR expression was significantly higher in the uninfected and infected group treated with dalbavancin respect to the other two groups (p<0.05), the wounds treated with dalbavancin showed the highest expression.

MMP-9 and MMP-1 expression was high in the untreated infected wounds, while a significant reduced expression was found in the other groups (p<0.05), in particular the treated wounds showed the lowest values.

### Analysis by Western blot of the Expression of EGFR, VEGF, MMP-1and MMP-9

The expression of EGFR, VEGF, MMP-1 and MMP-9 was tested by Western blot followed by densitometric analysis (Fig. 5). All proteins were detectable in uninfected, untreated infected, infected treated with dalbavancin and infected treated with vancomycin tissue samples. The expression of EGFR and VEGF was significantly higher in infected treated with dalbavancin tissue than all the other tissue samples (p < 0.05, Fig. 5). VEGF protein expression level was significantly lower in the untreated infected compared with uninfected and treated infected (p < 0.05 Fig. 5). MMP-1 and MMP-9 protein expression levels were significantly higher in the untreated infected compared with the other groups; in particular, the treatments with dalbavancin and Vamcomycin resulted in a significant decrease of protein expression levels (p < 0.05 Fig. 5).

**Figure 5.**
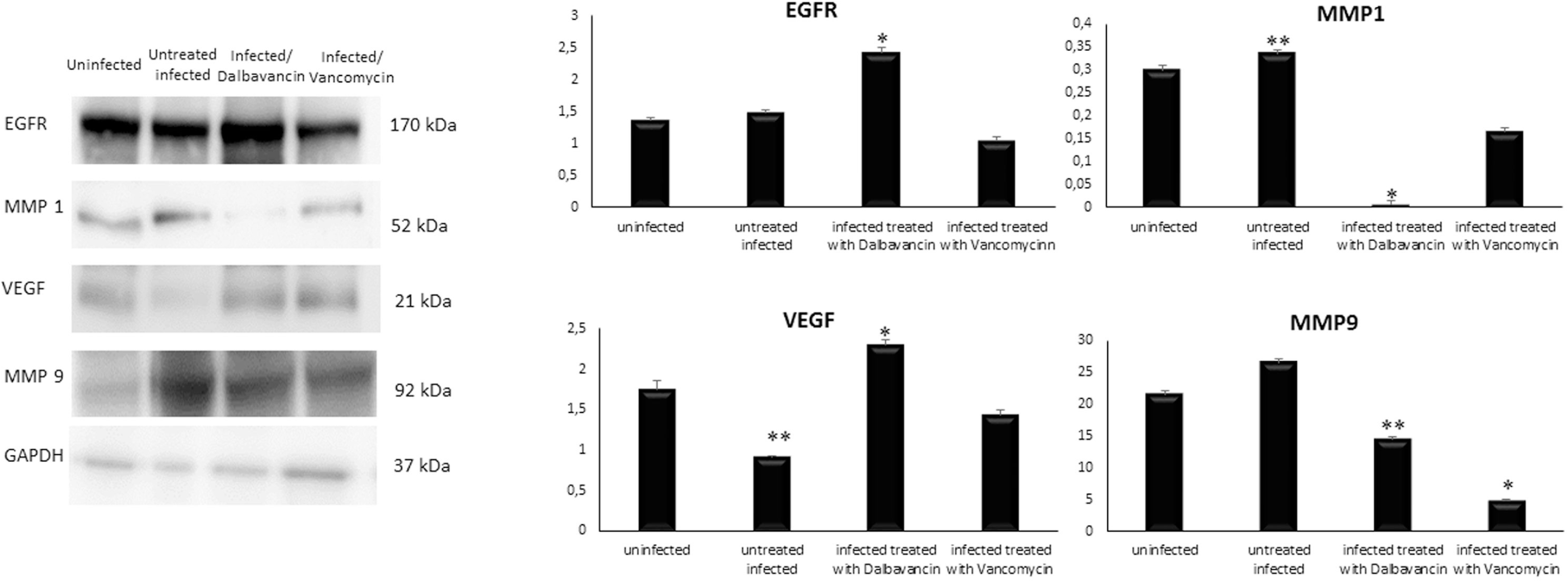
Analysis of the expression of EGFR, MMP-1, MMP-9 and VEGF. Western blot and densitometric analyses of EGFR, MMP-1, MMP-9 and VEGF, in a representative of the three performed experiments. Densitometric analyses of the immunoreactive bands is quantified as ratio between band relative to EGFR, VEGF, MMP-1, MMP-9 and, and GAPDH in corresponding samples, revealed by arbitrary units. Data are expressed as a mean ± SD from analyses performed on uninfected, untreated infected, infected treated with vancomycin and infected treated with dalbavancin tissue samples. * infected treated with dalbavancin versus uninfected, untreated infected, infected treated with vancomycin p < 0.05. ** untreated infected versus uninfected, infected treated with dalbavancin, infected treated with vancomycin p < 0.05.

## DISCUSSION

MRSA has demnostrated to be prevalent in ABSSSIs and in SSIs. Indeed, the rates of MRSA infection are increasing dramatically and MRSA has emerged as the most common cause of surgical infections (18).

Vancomycin is the first-choice antibiotic for the treatment of MRSA infection (19), but several papers described infections sustained by vancomycin-intermediate *S. aureus* (VISA) and heterogeneous VISA (hVISA) strains. Different studies reported that the host immune system has difficulty in eradicating VISA/hVISA, leading to chronic, recurrent, and persistent infections (20). Dalbavancin is an optimal alternative to conventional antibacterials for the treatment of ABSSSIs. Like other lipoglycopeptides, dalbavancin binds the *C*-terminal *acyl*-d-Ala-d-Ala subunit of peptidoglycan precursors. Furthermore, the positively charged *C*-terminal dimethylaminopropyl group may interact with the negative phospholipid head groups of the bacterial membrane (21). The fatty acyl group allows nonspecific protein binding that likely contributes to dalbavancin’s long plasma half-life and its ability to bind its target sites (21). This particular structure confers exceptional pharmacokinetic properties to dalbavancin and allow the administration in a two-dose regimen (9).

Dalbavancin demonstrated superior *in vitro* activity versus comparators, including vancomycin, with dramatically lower MICs without evidence of resistance through direct selection or serial passage (22,9).

Our microbiological results confirmed that both dalbanvancin and vancomycin are effective in reducing the bacterial load in wounds. These results are not surprising, considering that efficacy of these antibiotics was proven by the large clinical use and the numerous reports (23–25). Moreover, due to the long biological half-life, daily treatment didn’t result in better reduction of bacteria, as also previously demonstrated by other authors (26).

But, till now there is no published data implicating the effect of dalbavancin in wound healing comparing to vancomycin.

After external skin injury, such as chirurgical intervention, a rapid repair is needed to avoid dehydration and blood loss. Thus, a wound healing process involves a coordinated cascade of events comprising inflammation, formation of granulation tissue and dermis maturation (27–28). It has been shown that the bacteria are specifically able to alter stromal components and inflammatory cells and their responses. Infact, Human stromal cells demonstrated, to be susceptible to soluble factors of *S. aureus*, biofilms, decreasing cell differentiation, viability, migration and angiogenesis (29). Therefore *S. aureus* showed to have a dramatic effect on keratinocyte proliferation rates as well as an impedance to the migration of the keratinocytes, evidenced by delayed wound closure (30). Thus, we studied the effect of dalbanvancin *vs* vancomycin on the infected MRSA wound healing process through morphological studies besides an immunoistochemical evidences; moreover, to confirm our results we performed a western blot analysis.

As expected, in the infected but untreated groups, wound healing was still incomplete, with partial regeneration of the epidermis, poor organization of dermis layers, immature granulation tissue, and abundant inflammatory infiltrate. Both treatments showed to be efficacy on wound healing process. In particular, in dalbavancin treated group we observed a robust epidermal coverage with a reconstitution of the regular and keratinized epidermal lining, while epidermis in vancomycin treated group was still hypertrophic, revealing that epithelial healing problem is still occurring. Besides, in the present study we discovered elevated EGFR expression in the treated groups, although it was higher in dalbavancin treated mice. These immunohistochemical results correlated with the histological features and suggested that dalbavancin and vancomycin were able to induce EGFR expression and consequently to have an effect on wound healing. In particular, dalbavancin seemed to be more effective than vancomycin in inducing reepithelialization.

Epidermal keratinocytes are a rich source of EGFR ligands, and EGFR signaling has a major effect on the proliferation and differentiation of keratinocytes. Therefore, EGFR plays a key role in skin development and homeostasis (31) and its expression in the epithelium is a good indicator of the healing potential as observed in previuos studies (32,14). In our study MMP-1 and MMP-9 were decreased in uninfected and in both treated tissues, when compared with untreatd infected wounds. It has demonstrated that these metalloproteinases play a pivotal role, with their inhibitors, in regulating extracellular matrix (ECM) degradation and deposition that is essential for wound reepithelialization (33).

While, in normal tissue, MMP*s* are expressed at basal levels, when tissue remodeling is required (as in wound healing), they can be rapidly expressed and activated. Therefore, MMP*s* are upregulated during wound healing in the very early stages in epidermal cells, dermal cells, fibroblasts and blood cells (12).

Infact, it has been shown that MMP-1 expression peaks at day 1 after wounding in migrating basal keratinocytes at the wound edge followed by a gradual decrease until re-epithelialization is complete. Downregulation of MMP-1 seems to be important for normal tissue remodeling as high levels of MMP-1 in chronic nonhealing wounds are described (33). Similarly migrating keratinocytes at the leading edges secrete MMP-9, that modulate keratinocyte motility by degradation of proteins involved in cell–cell and cell–matrix adhesion, allowing re-epithelialization to occur (33). In particular, MMP-9 knockout (KO) mice, and MMP-9-deficient mice showed delayed wound closure with inhibition of cell proliferation (34–35), whereas the persistence of high level of MMP-9 has been associated with poor healing in several studies (36–38). These findings highlighted the role of MMP-9 in wound healing. In addition, when the balance of MMPs activitiy and ihnibition is disturbed the wound progress to a state of increased ECM degradation, disrupting the equilibrium of synthesis and remodeling of ECM components, determinant for succesful skin repaired (39). It is well known that *MMP*s play also a significant role in stimulation of angiogenesis in the proximity of wounds to accelerate recovery (33). In particular, the role of MMP-9 in angiogenesis include VEGF release (40–41). Actually, VEGF and MMP seem to regulate each other, contributing to wound repair (42). Previous studies have evidenced that keratinocytes express elevated VEGF as early as 1 day after injury and eventually in those cells that migrate to cover the defect. Epidermal labeling for VEGF mRNA reaches a peak after 2–3 days, and levels remain elevated during the granulation tissue formation and until epidermal coverage is complete (43). In this period, VEGF is upregulated to promote the angiogenesis (i.e., vascular dilation, permeability, migration, and proliferation), providing a conduit for nutrients and other mediators of the healing response (43). In our study we observed that VEGF expression was highly expressed in the uninfected and both treated infected groups, although the wounds treated with dalbavancin showed the highest VEGF values, underlining the positive effect of dalbavancin on angiogenesis.

In conclusion it can be speculated that both treatments may have positive effect not only on the bacterial load, but also on tissue healing after surgical infected skin wound by downregulation of MMP-1 and MMP-9 and increase of EGFR and VEGF expression. In particular this study showed faster healing with dalbavancin treatment that might be associated with a higher EGFR and VEGF levels. However, further studies are needed to clarify the additional effects of Dalbavacin on wound healing.

## MATERIALS AND METHODS

### Ethics

*In vivo* experiments were approved by the Institutional Animal Care Committee of the Ministery of Health and by the Animal Research Ethics Committee of IRCCS-INRCA (Istituto di Ricovero e Cura a Carattere Scientifico – Istituto nazionale di Riposo e Cura per Anziani) 767/2016 Pr 28/07/2016.

### Bacterial strains and drugs

*S. aureus* ATCC 43300, the reference MRSA strain, was used in the mouse infection model. Vancomycin and dalbavancin were diluted in accordance with manufacturers’ recommendations yielding 10 mg/mL stock solutions. Solutions were made fresh on the day of assay or stored at −80°C in the dark for short periods.

### Mouse infection model

6-month-old BALB/c mice weighting 28–30 g from the Specific Pathogen Free (SPF) animal facility of INRCA (Italian National Centre on Health and Science on Aging, Ancona, Italy) were used for all the studies. All the procedures involving animals were conducted in conformity with the institutional guidelines in compliance with the national (Legislative Decree n. 26, March 4, 2014; Authorization n.767/2016-PR issued July 28, 2016, by the Italian MoH) and international law and policies (EEC Council Directive 2010/63/EU). Mice were individually caged in visual, auditory and olfactory contact with other mice of the same experimental group (within ventilated cabinets cage systems) as well as in presence of environmental enrichments consisting of nesting materials and wooden toys. Mice were kept in a 12 h light dark cycle with food and drinking water ad libitum and allowed to equilibrate in the phenotyping area allocated in the Scientific and Technological Pole of INRCA for approximately 1 month before starting the experiments. After surgical intervention, all mice were housed individually under constant temperature (22 ± 2 °C) and humidity. The animals were fed with standard pellet food (Harlan Laboratories, Udine, Italy), and fresh daily tap water.

The study included a total of 48 animals divided into 4 groups (each composed of 12 mice): uninfected group (C0, sham control); infected but not treated group (C1); vancomycin group (infected and daily treated with vancomycin; C2); dalbavancin group (infected and weekly treated with dalbavancin; C3).

The MRSA ATCC43300 were grown in brain hearth infusion and diluted in saline to a final concentration of 5×10^7^ CFU/ml, prepared freshly at time of intervention.

Mice were anesthetized by an intramuscular injection of ketamine (50 mg/kg of body weight) and xylazine (8mg/kg of body weight), the hair on their back was shaved and the skin cleansed with 10% povidone-iodine solution (no animals dropped out due to infection or anesthetics). One full thickness wound was established through the panniculus carnosus on the back subcutaneous tissue of each animal. Small gauze was placed over each wound and then inoculated with 200 µl of bacterial culture previously diluted; in the control group the gauze was soaked only with sterile saline solution. The pocket was closed by means of skin clips. This procedure resulted in a local abscess at 24 h. One wound was created per animal. The animals were returned to individual cages and thoroughly examined daily. After 24 h, the wound was opened and washed with saline, the gauze was removed, and treatment started (44). The group C0 did not received any treatment. The groups C1, C2 were treated with daily intraperitoneal administration of 200 μl of saline and intraperitoneal vancomycin (10mg/kg) respectively. Group C3 was treated with intraperitoneal injection of 200 µl of dalbavancin at day 1 (20 mg/kg) and day 8 (10 mg/kg), mimicking human regimens (45). After 14 days, animals, were euthanized and a 1×2 cm area of skin, including the wound, was excised aseptically for histological and western blot examination (see below) and for bacterial count. For the bacterial analysis the samples were weighted and then homogenized in 1 ml phosphate-buffered saline (PBS) using a stomacher. Quantitation of viable bacteria was performed by culturing serial dilutions of the bacterial suspension on mannitol-salt agar plates at 37°C for 24-48 hours. The limit of detection for this method was approximately 10 CFU/ml.

### Histological and immunohistochemical staining

To understand the efficacy of the treatments in promoting wound healing, the re-epithelialized tissue of each group was investigated histologically. Five micrometer thick sections were stained with hematoxylin and eosin (H&E) according to standard protocol. The stained sections were observed under the light microscope (Nikon DS-Vi1, Nikon Instruments, EuropeBV, Kingston, Surrey, England) and scored according to wound repair fivetiered grading system based on epithelial presence, degree of stratification, degree of differentiation as well as maturational features of the granulation tissue, dermal tissue and presence of inflammatory cells (46) (Table 1).

In addition, identification of VEGF, MMP-1, MMP-9 and EGFR expression was carried out using immunohistochemistry of histological samples with a heat-mediated antigen retrieval process.

Briefly, 5 μm paraffin sections were deparaffinized and rehydrated by sequential immersion in xylene, ethanol, and water. Antigen retrieval was carried out by heating the sections in 0.1 M citrate buffer solution (pH 6.0) at 98 °C for 5 min via microwave. Then they were incubated with monoclonal anti-mouse VEGF (clone C-1; Santa Cruz Biotechnologies, Santa Cruz, CA), MMP-1 (clone H-300; Santa Cruz Biotechnologies, Santa Cruz, CA), MMP-9 (clone 2C3; Santa Cruz Biotechnologies, Santa Cruz, CA) and EGFR (clone 1005; Santa Cruz Biotechnologies, Santa Cruz, CA) at 4 °C overnight and immunostained using the streptavidin-biotin peroxidase technique (Envision universal peroxidase kit; Dako Cytomation, Milan, Italy). After incubation, the tissue sections were colored with 3,3-diaminobenzidine (DAB), stained with hematoxylin, and coverslipped with Eukitt mounting medium (Electron Microscopy Sciences, PA, USA).

Two investigators (G.L., A.Z.) blinded to the mice outcome, performed all counts separately. We counted the number of marker positive cells by using a light microscope (Nikon Eclipse E 600, Nikon Instruments, Europe B.V., Amsterdam, Netherlands) at x200 magnification in at least ten fields/sample and the positive cells were expressed as mean value ± standard deviation (SD). Images were captured with a Nikon DS-Vi1 digital camera (Nikon Instruments) connected to a computer. The area of each field (0.07 mm^2^) was standardized using NIS Elements BR 3.22 imaging software (Nikon Instruments).

### Protein extraction and immunoblotting

Total proteins were extracted from wound tissue samples and immediately pulverized in liquid nitrogen. The pulverized skins were homogenized and lysed in ice-cold Radio immunoprecipitation assay buffer [150 mM NaCl, 10 mM Tris (pH 7.2), 0.1 % sodium dodecyl sulfate, 1.0 % Triton X-100, and 5 mM ethylenediaminetetraacetic acid (pH 8.0)] containing protease inhibitor cocktail (Roche Applied Science, Indianapolis, IN, USA) for 45 minutes. The lysates were centrifuged at 14 000 rpm for 30 minutes at 4°C. The supernatant was collected and the protein concentration was determined. Protein concentration was determined using Bradford reagent (Sigma-Aldrich). Total protein extracts (30 μg) were reduced in dithiothreitol (DDT, 0.5 M) for 10 min at 70 °C and samples run on a 4–12 % gradient precast NuPAGE Bis–Tris polyacrylamide gel for 1 h at 200 V. After electrophoresis, the proteins in the gel were transferred to a nitrocellulose membrane, the blots were blocked with 5% nonfat dry milk in PBS containing 0.1% Tween-20 for 1 hours at room temperature. Membranes were incubating overnight with primary antibodies anti EGFR, VEGF, MMP-1 and MMP-9, (Santa Cruz Biotechnology, CA, USA) and GAPDH ((Santa Cruz Biotechnology) as endogenous control, followed by incubation with a secondary antibody conjugated to horseradish peroxidase (Santa Cruz Biotechnology). The detection of the proteins on the membrane was performed using the Clarity Western ECL Substrate kit (*Bio-rad*). The signals were captured using the Alliance Mini (*UVITEC Cambridge*) system. The UVITEC software carried out the quantization of the bands and the intensity of each band of interes) was normalized comparing it to the housekeeping GAPDH protein, used as loading control. Subsequently, the intensity of each tested band was compared to negative controls.

### Statistical analysis

All results were expressed as mean values ± SD. Differences between the groups were analyzed by one-way analysis of variance (ANOVA) tests and p values were determined using the Bonferroni test. Statistical analyses were performed using the SPSS 16 package (SPSS Inc., Chicago, IL, USA). Significance was set at p <0.05.

## Trasparency declaration

Nothing to declare.

## REFERENCES

1) Russo A, Concia E, Cristini F, De Rosa FG, Esposito S, Menichetti F, Petrosillo N, Tumbarello M, Venditti M, Viale P, Viscoli C, Bassetti M. 2016. Current and future trends in antibiotic therapy of acute bacterial skin and skin-structure infections. Clin Microbiol Infect 22 Suppl 2:S27–36

2) Ray GT, Suaya JA, Baxter R. 2013. Incidence, microbiology, and patient characteristics of skin and soft-tissue infections in a U.S. population: a retrospective population-based study. BMC Infect Dis 13:252.

3) Boucher HW, Corey GR. 2008. Epidemiology of methicillin-resistant *Staphylococcus aureus*. Clin Infect Dis 46:S344–S349.

4) Khan A, Wilson B, Gould IM. 2018. Current and future treatment options for community-associated MRSA infection. Expert Opin Pharmacother 19:457–470.

5) Holmes MA, Zadoks RN. 2011. Methicillin resistant *S. aureus* in human and bovine mastitis. J Mammary Gland Biol 16:373–382.

6) Pexara A, Solomakos N, Govaris A. 2013. Prevalence of methicillin-resistant *Staphylococcus aureus* in milk and dairy products. J Hell Vet Med Soc 64:17–34.

7) Ferreira JP, Anderson KL, Correa MT, Lyman R, Ruffin F, Reller LB, Fowler VG Jr. 2011.Transmission of MRSA between companion animals and infected human patients presenting to outpatient medical care facilities. PLoS One 6:e26978.

8) Barlam TF, Cosgrove SE, Abbo LM, MacDougall C, Schuetz AN, Septimus EJ, Srinivasan A, Dellit TH, Falck-Ytter YT, Fishman NO, Hamilton CW, Jenkins TC, Lipsett PA, Malani PN, May LS, Moran GJ, Neuhauser MM, Newland JG, Ohl CA, Samore MH, Seo SK, Trivedi KK. 2016. Implementing an Antibiotic Stewardship Program: guidelines by the infectious diseases society of America and the society for healthcare epidemiology of America. Clin Infect Dis 62(10):e51–77.

9) Boucher HW, Wilcox M, Talbot G, Puttagunta S, Das AF, Dunne MW. 2014. Once-weekly dalbavancin versus daily conventional therapy for skin infection. N Engl J Me 370 (23):2169–79

10) Tsoulas C, Nathwani D. 2015. Review of meta-analyses of vancomycin compared with new treatments for Gram-positive skin and soft-tissue infections: Are we any clearer? Int J Antimicrob Agents 46 (1):1–7.

11) Doma E., Rupp C., Baccarini M. 2013. EGFR-ras-raf signaling in epidermal stem cells: roles in hair follicle development, regeneration, tissue remodeling and epidermal cancers. Int J Mol Sci 14:19361–19384.

12) Gill SE, Parks WC. 2008. Metalloproteinases and their inhibitors: regulators of wound healing. Int J Biochem Cell Biol 40(6-7):1334–1347.

13) Amar S, Smith L, Fields GB. 2017. Matrix metalloproteinase collagenolysis in health and disease Biochim Biophys Acta Mol Cell Res 1864(11 Pt A):1940–1951.

14) Barrientos S, Stojadinovic O, Golinko MS, Brem H, Tomic-Canic M. 2008. Growth factors and cytokines in wound healing. Wound Repair Regen 16(5):585–601.

15) Werner S, Grose R. 2003. Regulation of wound healing by growth factors and cytokines. Physiol Rev 83(3):835–70.

16) Simonetti O, Cirioni O, Ghiselli R, Goteri G, Orlando F, Monfregola L, De Luca S, Zizzi A, Silvestri C, Veglia G, Giacometti A, Guerrieri M, Offidani A, Scaloni A. 2012. Antimicrobial properties of distinctin in an experimental model of MRSA-infected wounds. Eur J Clin Microbiol Infect Dis 31(11):3047–55.

17) Simonetti O, Cirioni O, Goteri G, Ghiselli R, Kamysz W, Kamysz E, Silvestri C, Orlando F, Barucca C, Scalise A, Saba V, Scalise G, Giacometti A, Offidani A. 2008. Temporin A is effective in MRSA-infected wounds through bactericidal activity and acceleration of wound repair in a murine model. Peptides 29(4):520–8.

18) Sganga G, Tascini C, Sozio E, Colizza S. 2017. Early recognition of methicillin-resistant *Staphylococcus aureus* surgicalsite infections using risk and protective factors identified by a group of Italian surgeons through Delphi method. World J Emerg Surg 12:25.

19) Liu C, Bayer A, Cosgrove SE, Daum RS, Daum RS, Fridkin SK, Gorwitz RJ, Kaplan SL, Karchmer AW, Levine DP, Murray BE, J Rybak M, Talan DA, Chambers HF. 2011. Clinical practice guidelines by the Infectious Diseases Society of America for the treatment of methicillin-resistant *Staphylococcus aureus* infections in adults and children. Clin Infect Dis 52(3):e18–e55.

20) Howden BP, Peleg AY, Stinear TP. 2014. The evolution of vancomycin intermediate *Staphylococcus aureus* (VISA) and heterogenous-VISA. Infect Genet Evol 575–582.

21) Economou N J, Nahoum V, Weeks SD, Grasty KC, Zentner IJ, Townsend TM, Bhuiya MW, Cocklin S, Loll PJ. 2012. A carrier protein strategy yields the structure of dalbavancin. A carrier protein strategy yields the structure of dalbavancin. J Am Chem Soc 134(10): 4637–464510

22) Ramdeen S, Boucher HW. 2015. Dalbavancin for the treatment of acute bacterial skin and skin structure infections. Expert Opin Pharmacother 16(13):2073–81.

23) Bassetti M, Peghin M, Castaldo N Giacobbe DR. 2019. The safety of treatment options for acute bacterial skin and skin structure infections. Expert Opin Drug Saf 18(8):635–650

24) Savoldi A, Azzini AM, Baur D, Tacconelli E. 2018. Is there still a role for vancomycin in skin and soft-tissue infections? Curr Opin Infect Dis 31(2):120–130

25) Mendes RE, Castanheira M, Farrell DJ, Flamm RK, Sader HS, Jones RN. 2016. Update on dalbavancin activity tested against Gram-positive clinical isolates responsible for documented skin and skin-structure infections in US and European hospitals (2011-13). J Antimicrob Chemother 71:276–8.

26) Malabarba A, Goldstein BP. 2005. Origin, structure, and activity in vitro and in vivo of dalbavancin. J Antimicrob Chemother 55 Suppl 2:ii15–20.

27) Chow O, Barbul A. 2014. Immunonutrition: role in wound healing and tissue regeneration. Ad. Wound Care 3:46–53.

28) Landén NX, Li D,Ståhle M. 2016. Transition from inflammation to proliferation: a critical step during wound healing. Cell Mol Life Sci 73: 3861–3885.

29) Ward CL, Sanchez CJ Jr, Pollot BE, Romano DR, Hardy SK, Becerra SC, Rathbone CR, Wenke JC. 2015. Soluble factors from biofilms of wound pathogens modulate human bone marrow-derived stromal cell differentiation, migration, angiogenesis, and cytokine secretion. BMC Microbiol 15: 75.

30) Chaney SB, Ganesh K, Mathew-Steiner S Stromberg P, Roy S, Sen CK, Wozniak DJ. 2017. Histopathological comparisons of Staphylococcus aureus and Pseudomonas aeruginosa experimental infected porcine burn wounds. Wound Repair Regen 25(3):541–549

31) Zhang Z, Xiao C, Gibson, M Bass SA, Khurana Hershey GK. 2014. EGFR signaling blunts allergen-induced IL-6 production and th17 responses in the skin and attenuates development and relapse of atopic dermatitis. J Immunol 192(3):859–866.

32) Simonetti O, Lucarini G, Orlando F, Pierpaoli E, Ghiselli R, Provinciali M, Castelli P, Guerrieri M, Di Primio R, Offidani A, Giacometti A, Cirioni O.2017. Role of Daptomycin on Burn Wound Healing in an Animal Methicillin-Resistant Staphylococcus aureus Infection Model. Antimicrob Agents Chemother 24;61, pii: e00606–17.

33) Caley MP, Martins VL, O’Toole EA. 2015. Metalloproteinases and wound healing. Adv Wound Care (New Rochelle) 4(4):225–234.

34) Hattori N, Mochizuki S, Kishi K, Nakajima T, Takaishi H, D’Armiento J, Okada Y. 2009. MMP-13 plays a role in keratinocyte migration, angiogenesis, and contraction in mouse skin wound healing. Am J Pathol 175:533–546.,

35) Mohan R, Chintala SK, Jung JC, Villar WV, McCabe F, Russo LA, Lee Y, McCarthy BE, Wollenberg KR, Jester JV, Wang M, Welgus HG, Shipley JM, Senior RM, Fini ME. 2002. Matrix metalloproteinase gelatinase B (MMP-9) coordinates and effects epithelial regeneration. J Biol Chem 277:2065–2072.

36) Lazaro JL, Izzo V, Meaume S Davies AH, Lobmann R, Uccioli L. 2016. Elevated levels of matrix metalloproteinases and chronic wound healing: an updated review of clinical evidence. J Wound Care 25(5):277–87.

37) Simonetti O, Cirioni O, Lucarini G Orlando F, Ghiselli R, Silvestri C, Brescini L, Rocchi M, Provinciali M, Guerrieri M, Di Primio R, Giacometti A, Offidani A. 2012. Tigecycline accelerates staphylococcal-infected burn wound healing through matrix metalloproteinase-9 modulation. J Antimicrob Chemother 67(1):191–201.

38) Simonetti O, Lucarini G, Cirioni O Zizzi A, Orlando F, Provinciali M, Di Primio R, Giacometti A, Offidani A. 2014. Delayed wound healing in aged skin rat models after thermal injury is associated with an increased MMP-9, K6 and CD44 expression. Erratum in: Burns. 40(1):175 related to Burns. 2013 Jun;39(4):776–87.

39) McCarty SM, Percival SL. 2013. Proteases and Delayed Wound Healing. ADV Wound Care (New Rochelle) 2(8):438–447.

40) Bergers G, Brekken RA, McMahon G, Vu TH, Itoh T, Tamaki K, Tanzawa K, Thorpe P, Itohara S, Werb Z, Hanahan D. 2000. Matrix metalloproteinase-9 triggers the angiogenic switch during carcinogenesis. Nat Cell Biol 2:737–44.

41) Ardi VC, Van den Steen PE, Opdenakker G Schweighofer B, Deryugina EI, Quigley JP. 2009. Neutrophil MMP-9 proenzyme, unencumbered by TIMP-1, undergoes efficient activation *in vivo* and catalytically induces angiogenesis via a basic fibroblast growth factor (FGF-2)/FGFR-2 pathway. J Biol Chem 284:25854–66.

42) Howard EW, Soo C, Hunter S. Zhang X, Shaw WW, Longaker MT. 1997. Differences in MMP and TIMP gene expression during wound repair. Surg Forum 48:688–91.

43) Bao P, Kodra A, Tomic-Canic M, Golinko MS, Ehrlich HP, Brem H. 2009. The role of vascular endothelial growth factor in wound healing. J Surg Res 153(2):347–58.

44) Silvestri C, Cirioni O, Arzeni D, Ghiselli R, Simonetti O, Orlando F, Ganzetti G, Staffolani S, Brescini L, Provinciali M, Offidani A, Guerrieri M, Giacometti A. *In vitro* activity and *in vivo* efficacy of tigecycline alone and in combination with daptomycin and rifampin against Gram-positive cocci isolated from surgical wound infection. Eur J Clin Microbiol Infect Dis 31(8):1759–64.

45) Seltzer E, Dorr MB, Goldstein BP, Perry M, Dowell JA, Henkel T; Dalbavancin Skin and Soft-Tissue Infection Study Group. 2003. Dalbavancin Skin and Soft-Tissue Infection Study Group.Once-weekly dalbavancin versus standard-of-care antimicrobial regimens for treatment of skin and soft-tissue infections.Clin Infect Dis 37(10):1298–303.

46) Simonetti O, Cirioni O, Cacciatore, Baldassarre L, Orlando F, Pierpaoli E, Lucarini G, Orsetti E, Provinciali M, Fornasari E, Di Stefano A, Giacometti A, Offidani A.2016. Efficacy of the Quorum Sensing Inhibitor FS10 Alone and in Combination with Tigecycline in an Animal Model of Staphylococcal Infected Wound. PLoS One 11(6):e0151956.

